# Various plasmid strategies limit the effect of bacterial Restriction-Modification systems against conjugation

**DOI:** 10.1101/2024.06.17.599295

**Authors:** Tatiana Dimitriu, Mark D. Szczelkun, Edze R. Westra

## Abstract

In bacteria, genes conferring antibiotic resistance are mostly carried on conjugative plasmids, mobile genetic elements which spread horizontally between bacterial hosts. Bacteria carry defence systems which defend them against genetic parasites, but how effective these are against plasmid conjugation is poorly understood. Here, we study to what extent Restriction-Modification (RM) systems – by far the most prevalent bacterial defence systems - act as a barrier against plasmids. Using 10 different RM systems and 13 natural plasmids conferring antibiotic resistance in *Escherichia coli*, we uncovered variation in defence efficiency ranging from none to 10^5^-fold protection. Further analysis revealed genetic features of plasmids that explain the observed variation in defence levels. First, the number of RM recognition sites present on the plasmids generally correlates with defence levels, with higher numbers of sites being associated with stronger defence. Secondly, some plasmids encode methylases that protect against restriction activity. Finally, we show that a high number of plasmids in our collection encode anti-restriction genes that provide protection against several types of RM systems. Overall, our results show that it is common for plasmids to encode anti-RM strategies, and that, as a consequence, RM systems form only a weak barrier for plasmid transfer by conjugation.

## Introduction

Plasmids are extra-chromosomal DNA molecules which replicate independently from the bacterial chromosome; conjugative plasmids can transmit horizontally between bacterial cells as they encode a conjugation machinery allowing transfer between adjacent cells (1). Conjugative plasmids are central to bacterial ecology and evolution because they frequently confer adaptive traits to bacteria, such as metal resistance, antimicrobial resistance (AMR), or virulence traits (2). In particular, conjugative plasmids play a major role in the spread of AMR genes among commensal and pathogenic bacterial species (3). In *Enterobacteriaceae*, a few major conjugative plasmid families are responsible for AMR including resistance to clinically relevant antibiotics (4, 5). It is thus key to understand what shapes conjugative plasmid transmission among bacteria.

Plasmid population dynamics is primarily controlled by two parameters: the effect of a plasmid on its bacterial host’s fitness (which impacts plasmid vertical transmission), and its rate of horizontal transmission between cells by conjugation (6). There is now an extensive literature on plasmid fitness effects and their molecular determinants across plasmids and hosts (7), whereas variation in conjugation rates is less well understood, despite spanning many orders of magnitude (8). Conjugation will impact not only a plasmid’s ability to persist within a population (9), but also plasmid dissemination at larger scales, driving epidemics within species (10) and AMR gene dissemination across species (11). Conjugation rates themselves are determined by multiple factors: expression of the plasmid’s conjugative machinery, regulated by plasmid-encoded genes as well as the donor regulatory network (1, 12); but also successful mating pair formation and mating pair stabilisation, which can depend on the expression of specific receptors on the recipient cell’s surface (13); and finally successful establishment of plasmids in the recipient cell. For this last step to be successful, plasmids must overcome any defence systems present in the recipient.

Bacterial defence systems defend bacteria against molecular parasites, and their activity is mostly studied against bacterial viruses, or phages (14). However the most prevalent defence systems, Restriction-Modification (RM) systems and CRISPR-Cas systems, both act by recognizing specific DNA sequences, and as such can target plasmids as well as phages (15). At evolutionary timescales, these defence systems have been shown to impact horizontal gene transfer (16, 17), and specifically the distribution of AMR mobile elements in bacterial pathogens (18–20). RM systems are by far the most common defences in bacteria, with on average more than two RM systems per bacterial genome (14). These innate immune systems use DNA modification to distinguish self from non-self. Specifically, they encode a restriction endonuclease, which cleaves DNA following the recognition of an unmodified recognition site, and a methyltransferase which protects cellular DNA by methylating the recognition sites. Incoming DNA with unmodified recognition sites is recognized as non-self, and restricted.

Restriction endonucleases can cleave any double-stranded DNA molecules carrying unmodified recognition sequences. Recognition sites are commonly present in several copies on any DNA sequences longer than a few kilobases, which includes conjugative plasmids. Indeed, restriction was shown in the early days of molecular biology to have an effect against plasmids, and specifically against DNA entry into cells by conjugation (21). Yet, restriction efficiency is often found to be low against conjugation (22), especially when compared to other gene transfer mechanisms (23). For instance, when first investigated, EcoKI restriction was observed to restrict phage λ by more than 10^5^-fold, but conjugation by only 100-fold (24). Various causes have been suggested for this relative resistance to restriction, including the fact that conjugating plasmids enter the recipient as single-stranded DNA before being replicated (25), avoidance of recognition sites in some plasmid sequences (26), and carriage of anti-restriction and protective functions by plasmids (27). For a few model plasmids, recognition site avoidance and carriage of specific genes has been shown experimentally to provide protection against restriction in the context of conjugation (28–30). However, the experimental data is scarce, limiting our ability to infer general causal relationships between the presence of RM systems and plasmid conjugation rates, and consequently our understanding of this relationship is lagging that of RM-phage interactions (31, 32).

Here, we provide a more systematic analysis of how efficient different classes of RM systems are against AMR plasmid conjugation. To study plasmids representative of the known diversity of plasmid types, with a focus on AMR plasmids in bacterial pathogens, we assembled a collection of 13 AMR plasmids, which include well-studied model plasmids and plasmids isolated from clinical environments (9, 33, 34), and which belong to 9 different plasmid families playing a key role in AMR transmission (5, 35). These plasmids vary in several key aspects including size, replicon and conjugative transfer type (Table S1). We measure conjugation of these plasmids towards a collection of recipient strains carrying 10 different restriction systems all native to *Escherichia coli* (36). These systems belong to RM Types I, II and III, the three types of classical RM systems differing in molecular structure and mechanistic properties (37). Type I RM are multiprotein complexes, with cleavage requiring the interaction of two endonuclease subunits, and happening at a distance from the recognition site. Type II RM cleave within or close to their recognition site. Finally, type III RMs are multiprotein complexes with an asymmetric recognition site, and cleavage requires two recognition sites on the same DNA in inverted repeat. We use four Type I systems and four Type II systems, with several subtypes of each, and two Type III systems, mirroring the relative abundance of these different Types across prokaryotic genomes (38). Finally, we explore potential causes for the observed variation in restriction efficiency.

## Methods

### Strains, plasmids, and growth conditions

Bacterial strains were grown at 37°C in Luria-Bertani (LB) broth, with agitation at 180 rpm. The 13 wildtype conjugative plasmids used in this study are listed in Table S1 with relevant characteristics: they vary in size, replicon, MOB (mobilisation) and MPF (mating pair formation) types, as well as AMR gene content and antibiotics used for selection (Table S1). For selection of plasmid-carrying strains, antibiotics were used at the following concentrations: trimethoprim 25 mg/L, streptomycin 100 mg/L, tetracycline 10 mg/L or ampicillin 100 mg/L. Individual gene deletion mutants of these conjugative plasmids were also used and described in the next section.

The 10 *Escherichia coli* RM systems we used (36) are listed in Table S2 with their recognition sequence: they vary in mechanistic type (four Type I systems, four Type II systems and two type III systems). Type I systems are encoded on the chromosome, Type II and Type III systems are encoded on plasmids maintained with ampicillin (100 mg/L) and chloramphenicol (25 mg/L), respectively.

Conjugation assays used *E. coli* MG1655 ΔdapA::ErmR ΔhsdS::KnR or MG1655 ΔdapA::ErmR ΔhsdS::frt Δdcm::frt (abbreviated as RM0 Δdap). ΔdapA strains cultures were supplemented with diaminopimelic acid (DAP) 300 µM.

### Cloning

*E. coli* MG1655 ΔdapA::ErmR and MG1655 ΔdapA::ErmR ΔhsdS::KnR were generated by P1 transduction of the ΔdapA::ErmR locus from MFDpir (39) to MG1655 and MG1655 ΔhsdS::KnR (40), using selection with erythromycin 400mg/L + DAP 300µM + citrate 5mM. To obtain RM0 Δdap, the KnR kanamycin resistance cassette was first excised from MG1655 ΔhsdS::KnR using pCP20 (41), then the *dcm* gene was inactivated by insertion of *cat* from pKD3 plasmid (41), and the *cat* cassette was excised using pCP20. Finally, the ΔdapA::ErmR locus was transduced from MFDpir as described above.

Plasmid gene knock-outs were performed using λred recombination (41): pKD3’s chloramphenicol resistance cassette was amplified using primers including a tail with 50-bp homologies to regions adjacent to the target genes, using Q5 High-Fidelity DNA Polymerase (New England BioLabs). Diagnostic PCRs were performed using Thermo Dream Taq^TM^ PCR. Recombineering and diagnostic primers are listed in Table S2.

Type II and Type III RM system plasmids were transformed into MG1655 ΔhsdS::KnR (40); conjugative plasmids were moved to the donor strains by conjugation using selection with erythromycin 300 mg/L + DAP 300 µM.

### Conjugation assays and measure of restriction efficiency

Plasmid-carrying donors and recipient strains were first grown overnight in glass vials, without antibiotic selection for conjugative plasmids. Conjugation assays were run in a total volume of 1 mL of LB broth supplemented with DAP 120 µM in 24-well plates: for each conjugation, 50 μL of donor culture and 50 μL of recipient culture were added to pre-warmed medium and incubated at 37°C with shaking at 180 rpm. Each donor x recipient combination was run in 4 or 5 replicates depending on the experiment, and control assays with donor only and recipient only were run in each experiment. After 3 to 4 hours, the conjugation mixes were serially diluted in M9 salts and droplets of appropriate dilutions were plated on LB-agar + antibiotic + DAP, LB-agar and LB-agar + antibiotic to estimate densities of donors, recipients, and transconjugants respectively. As R6K plasmid had low transfer efficiency, for this plasmid the whole conjugation mix was also plated without serial dilution in order to improve the detection threshold. The antibiotics used for selection of plasmid-bearing cells are shown in Table S1 for each plasmid. For pCF12, pOXA-48 and pCT plasmids, restriction by Type II RM systems was not measured as these plasmids carry only β-lactamase genes as selective markers, and the Type II RM systems are located on plasmids which also confer resistance to β-lactams.

Transfer efficiency was estimated as **γ** (mL/cell/h) = T / DRt, where T, D and R respectively indicate the cell density of transconjugants, donors and recipients (cells.mL^-1^), and t is the incubation time (h). When no transconjugant colony was present, a threshold transfer efficiency was calculated by assuming that 0.5 transconjugant colony was detected. Restriction efficiency for a given plasmid and RM system was calculated as RE = **γ**^RM+^ / **γ**^RM-^ where **γ**^RM+^ is the transfer efficiency towards the recipient strain carrying the RM system and **γ**^RM+^ is the transfer efficiency from the same donor towards MG1655 ΔhsdS::KnR. To calculate restriction efficiency, replicates were paired arbitrarily by replicate number.

### Sequencing and bioinformatics analyses

Two conjugative plasmids for which a sequence was not available publicly at the time of study, RIP113 and pCU1, were sequenced via MicrobesNG (http://www.microbesng.uk) using hybrid whole-genome sequencing, and deposited in the European Nucleotide Archive, project PRJEB76555. Plasmids MOB type, MPF type and replicon were determined using COPLA (42). The number of recognition sites on conjugative plasmids for each RM system was determined using Geneious Prime ^®^ 2023.2.1. In order to look for putative anti-restriction functions, we re-annotated all sequences using Bakta (43), then used both Bakta output as well as the original plasmid annotations.

### Statistical analyses

All statistical analyses were performed with R version 4.3.2 (44). For data processing and plotting we also used packages *reshape* (45) and *dplyr* (46), and packages *cowplot* (47), *ggpmisc* (48) and *broom* (49) to plot statistical parameters.

For each experiment, statistical parameters are reported in the figure legends or within the results section. Transfer efficiency and restriction efficiency (RE) were always log-transformed before statistical analysis. To analyse the effect of recognition site number across the whole dataset of wild-type plasmids (Fig. 2), a value of 0.5 was added to the data for two plasmid x RM combinations (EcoKI sites for RIP113, EcoVIII sites for pOXA-48) for which the number of recognition sites was zero.

**Figure 1:**
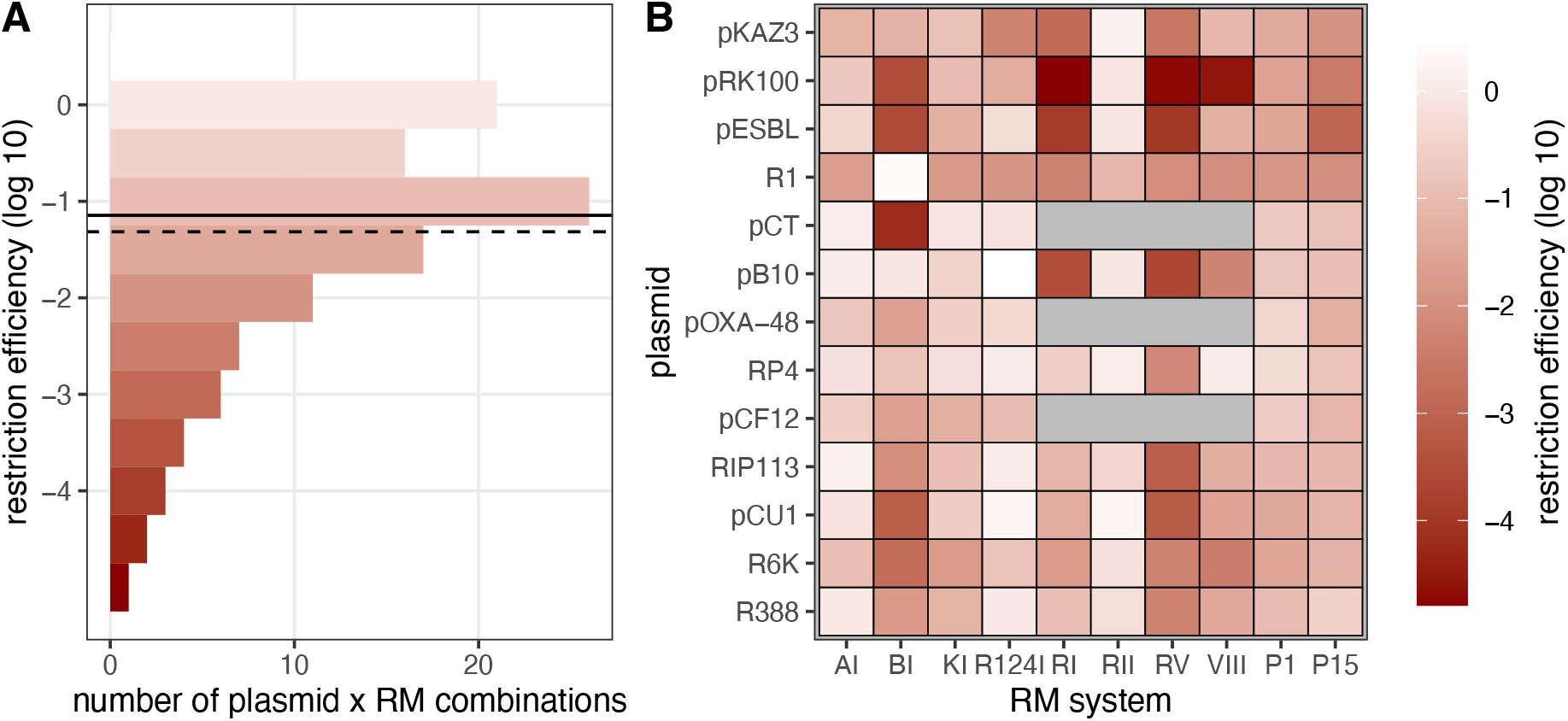
restriction efficiency of *E. coli* RM systems across wildtype AMR plasmids. In A, a histogram of the distribution of restriction efficiency (RE) across plasmid x RM combinations is shown. The solid and dashed lines indicate respectively median and mean RE. B shows average restriction efficiency per plasmid x RM system combination (geometric mean of n≥4 replicates) as a colour scale. Gray squares indicate non-measured combinations (see Methods).

**Figure 2:**
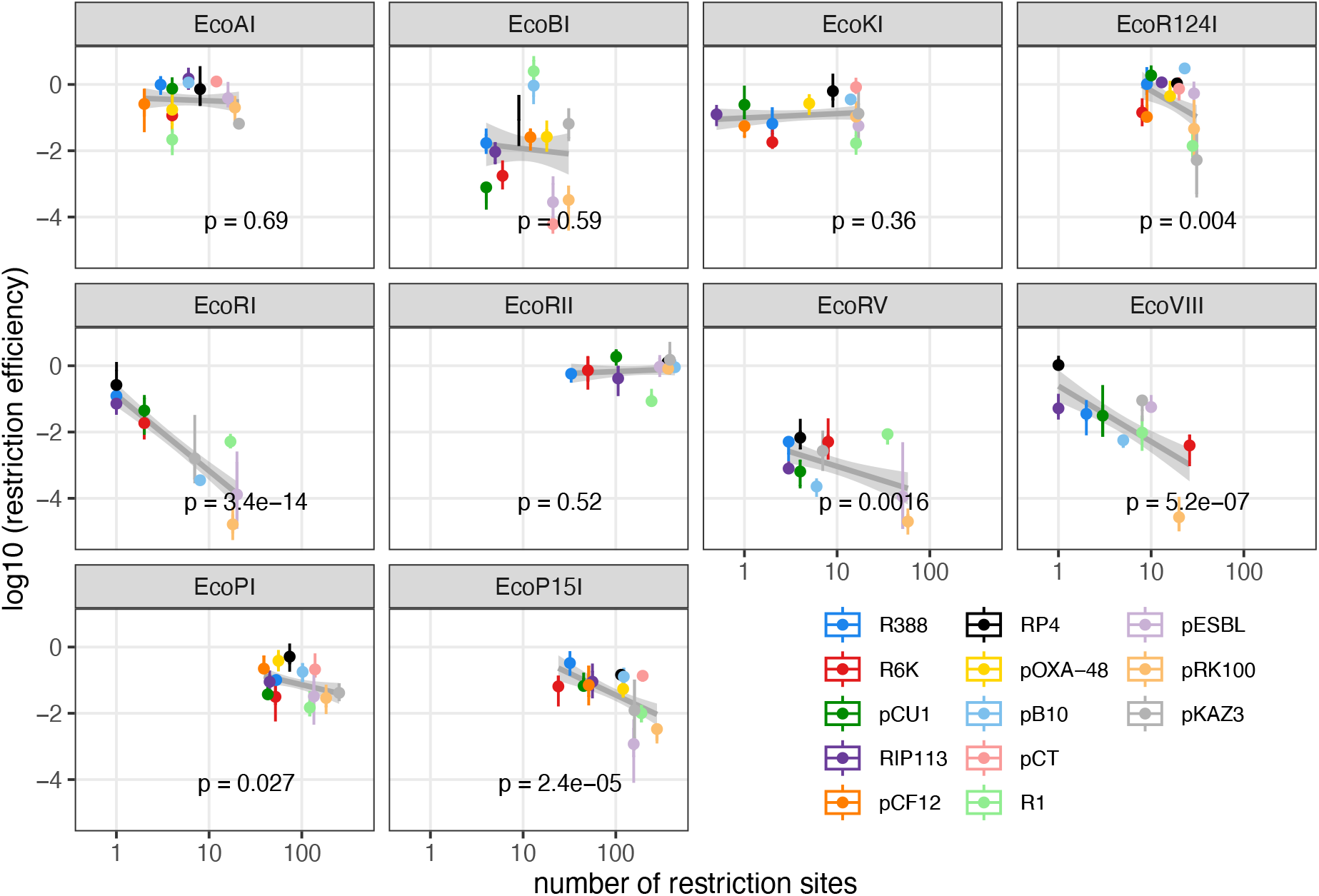
Effect of RM recognition site number on restriction efficiency across wildtype conjugative plasmids. Log-transformed RE is shown as a function of recognition site number for each RM system. Dots and vertical lines indicate respectively the average restriction efficiency per plasmid and the SEM (n≥4 replicates), with the colour indicating plasmid identity. The grey line represents the model fit on log_10_ (RE) ∼ log_10_ (n_sites_) for each system, and shaded areas show 95% confidence intervals, with p-values indicated. RM systems are ordered per mechanistic type (top row: type I, middle row: type II, bottom row: type III).

To analyse the effect of plasmid mutants on restriction, we first asked if plasmid mutants influenced restriction efficiency for specific RM systems with a linear model, RE ∼ mutant identity * RM system; in the case of pKAZ3 anti-restriction mutants (Fig. 4), the experiment was performed twice and ‘experiment’ was added as a factor (RE ∼ experiment + mutant identity * RM system). When the interaction term between mutant and RM system was significant (p<0.05), each mutant RE was compared to wild-type RE with an unpaired t-test, and we applied a Bonferroni correction to correct for multiple testing (10*n tests, where 10 is the number of RM systems and n is the number of mutants tested).

**Figure 3:**
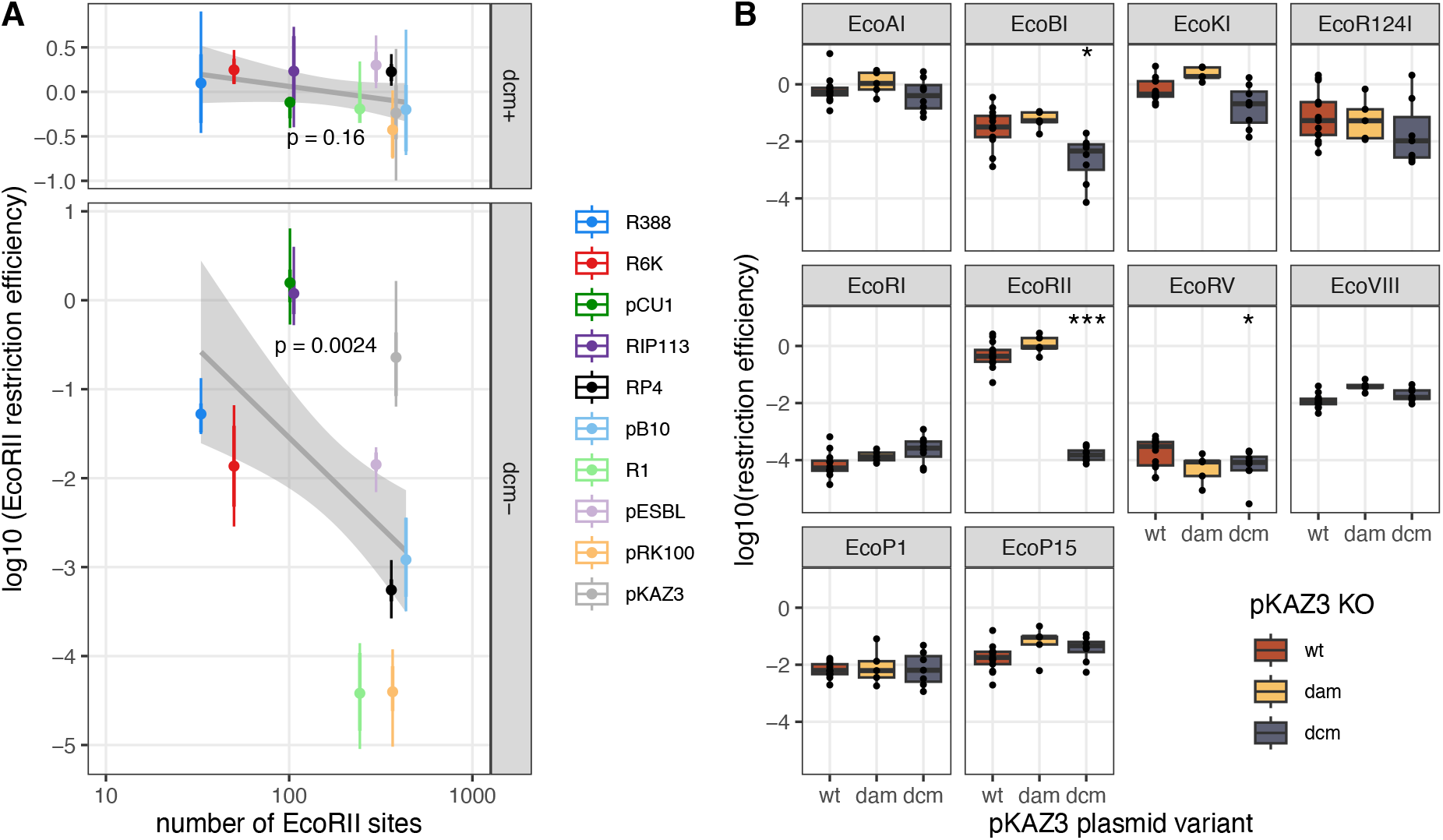
effect of chromosome and plasmid methylases on restriction efficiency. In A, log-transformed EcoRII RE is shown as a function of recognition site number on the conjugative plasmids, with the donor strain being *dcm+* (top) or *dcm-* (bottom). Dots and lines indicate respectively the average restriction efficiency per plasmid and the SEM (n = 4 replicates), with the colour indicating plasmid identity. The grey lines represent the model fit on log_10_ (RE) ∼ log_10_ (n_sites_), and shaded areas show 95% confidence intervals, with p-values indicated. In B, log-transformed RE for 10 *E. coli* RM systems is shown from a *dcm-* donor strain for wild-type pKAZ3, pKAZ3 Δ*dam* and pKAZ3 Δ*dcm*. The centre value of the boxplots shows the median, boxes the first and third quartile, and whiskers represent 1.5 times the interquartile range. Individual replicates are shown as dots (n≥4). Asterisks show RM treatments for which the mutant plasmid is significantly different from the wild-type plasmid after Bonferroni correction (*, 0.005 < *p* < 0.05; *** *p* < 0.0005).

**Figure 4:**
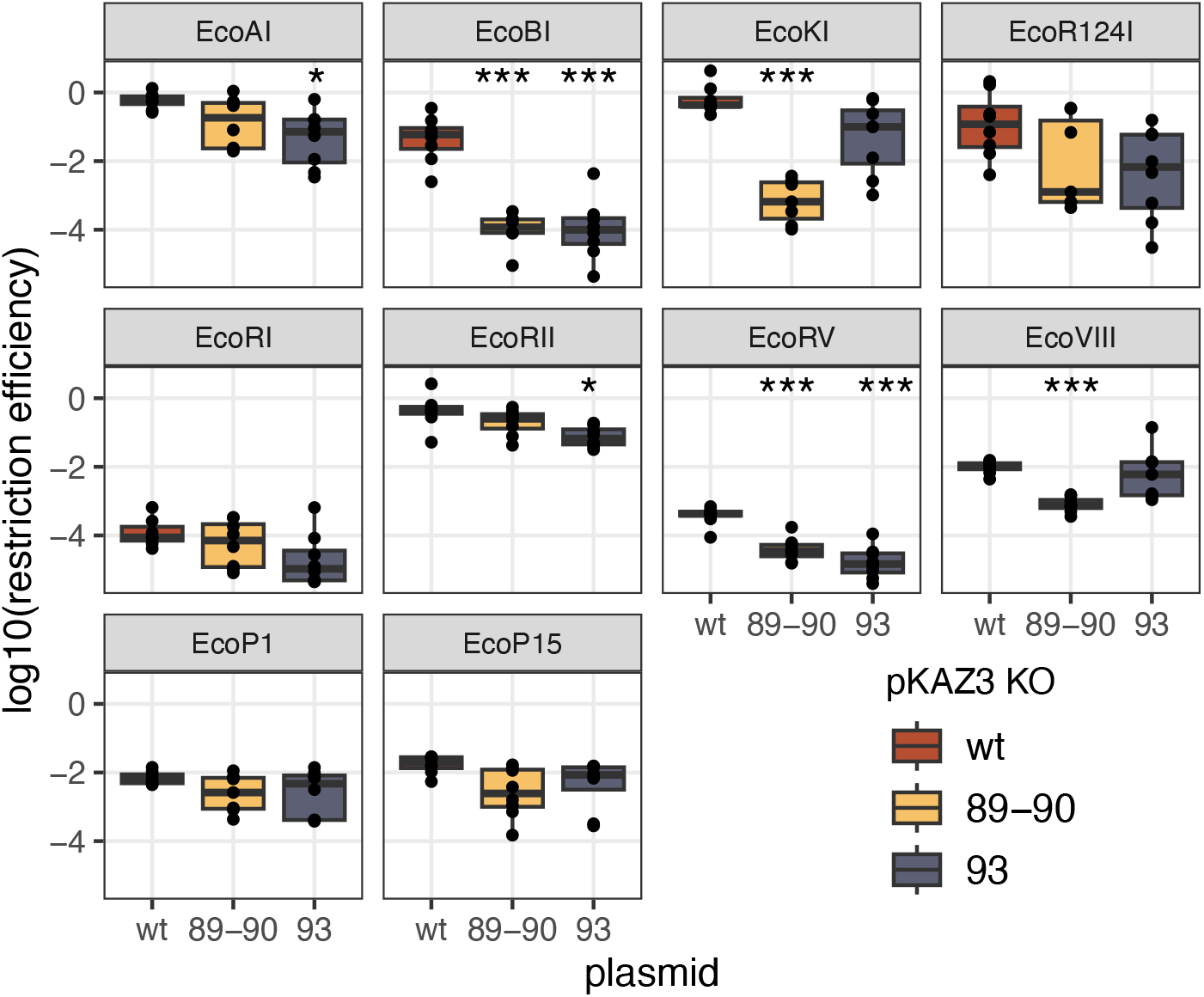
effect of plasmid anti-restriction functions on restriction efficiency. Log-transformed restriction efficiency for 10 *E. coli* RM systems is shown from a *dcm-* donor strain for wild-type pKAZ3, pKAZ3 Δ*vcrx089-90* and pKAZ3 Δ*vxrx093*. The centre value of the boxplots shows the median, boxes the first and third quartile, and whiskers represent 1.5 times the interquartile range. Individual replicates are shown as dots (n ≥ 4). Asterisks show RM treatments for which the mutant plasmid is significantly different from the wild-type plasmid after Bonferroni correction (*, 0.005 < *p* <0.05; *** *p* <0.0005).

## Results

### Restriction efficiency across *E. coli* conjugative plasmids and RM systems

To evaluate how effective restriction is against conjugative transmission, we first measured transfer efficiency for 13 wildtype plasmids in liquid culture, towards recipients carrying no or one of 10 *E. coli* RM systems. Transfer efficiency in the absence of RM systems varied between ∼ 10^-11^ and ∼ 10^-16^ mL/cell/h depending on the plasmids (Fig. S1). We then calculated restriction efficiency (RE) by comparing transfer efficiency for the same plasmid between recipients that do or do not carry a RM system.

RE values varied by 5 orders of magnitude depending on the plasmid and RM system (Fig. 1). Most combinations had relatively low RE, with a median effect of 14-fold and average effect of 20.7-fold (Fig. 1A). Overall, RE varied according to RM system identity, plasmid identity, but also their combination (Fig. 1B), with all these factors having a significant impact: in a linear model with log_10_ (RE) ∼ RM system * plasmid, RM system identity (F_9,449_ =211.9, p<2 e^-16^), plasmid identity (F_12,449_ =57.1, p<2.e^-16^) and their interaction (F_96,449_ =14.8, p<e^-16^) all had significant effects on RE. Type II EcoRII did not restrict plasmid conjugation at all, whereas Type II EcoRV had the strongest RE, with on average 1000-fold restriction across plasmids. When grouping RM systems by types, we found that Type II RM systems were significantly more efficient than Types I or III (log_10_ (RE) ∼ RM type * plasmid, interaction effect F_21,531_ =2.45, p=4.10^-4^ ; TukeyHSD test on RM type effect, Type II vs Type I 6.8-fold effect, p=0.000, Type III vs Type II, 3.5 fold effect, p=0.00003). Conjugative plasmids also varied in their average sensitivity to restriction: RP4 plasmid was restricted less than 3-fold on average, whereas pRK100 was restricted 290-fold on average. Yet, the strong effect of the interaction term, with some plasmids like pB10 displaying very variable sensitivity to restriction, suggests that specific factors determine a plasmid’s sensitivity to a given RM system.

### Effect of the number of RM recognition sites on RM efficiency

As RM systems are characterised by their sequence specificity, cleaving DNA after recognising specific unmethylated recognition sites (Table S2), we explored the effect of RM recognition site number for each plasmid on restriction efficiency (Fig. 2). Recognition site number varies across RM systems and plasmids (Fig. S2). Overall it is lower for Type I systems (median 12 per plasmid) and Type II systems (median 13), and highest for Type III systems (median 108).

We asked if the number of recognition sites has a statistically significant effect on RE for each RM system (RM). We included potential plasmid effects not due to their number of recognition sites (n_sites_) for a particular enzyme, using the linear model: log_10_ (RE) ∼ RM * log_10_ (n_sites_) + RM * plasmid. We found that both RM and n_sites_ had a highly significant effect (respectively F_9,449_ = 71, *p* < 2.10^-16^ and F_1, 449_ = 103, *p* < 2.10^-16^), and the interaction between RM and n_sites_ was also significant (F_9, 449_ = 8.99, *p* < 2.10^-16^). In addition, plasmid identity also had a significant effect after accounting for recognition site number (F_12, 449_ = 11.3, *p* < 2.10^-16^), as well as plasmid interaction with RM type (F_86,449_ = 13.4, *p* < 2.10^-16^).

Thus, an increased number of recognition sites overall increases restriction, but this varies depending on the specific RM system considered (Figure 2). When considering RM systems individually, n_sites_ has a significant effect for three of four Type II systems (EcoRI, EcoRV, and EcoVIII), with the remaining Type II system, EcoRII, having little restriction efficiency against any plasmids, independently of n_sites_ . n_sites_ effect was also significant for both Type III RM systems (EcoP1 and EcoP15), but with relatively low RE overall despite plasmids carrying more restriction sites than for Type II systems. Finally, n_sites_ had a significant effect for only one (EcoR124I) of the four Type I systems tested, with EcoAI, EcoBI and EcoKI being unaffected by n_sites_ despite strong variation in plasmid susceptibility to restriction (in particular, EcoBI RE varied by 4 orders of magnitude, but independently of n_sites_).

This absence of dependence of some RM systems on n_sites_, in particular Type I RM systems, and the significant effect of plasmid identity on RE, both suggest that additional plasmid traits must impact restriction efficiency. Next, we searched for putative anti-restriction functions carried by conjugative plasmids.

### Plasmid carriage of putative anti-restriction genes

The genomes of the 13 conjugative plasmids contain a variety of genes which annotations suggest anti-restriction activity (Table 1). First, many plasmids carry annotated DNA methylase genes: RIP113 plasmid carries the full EcoRII RM system, and all large plasmids (>90kb) carry at least one orphan methyltransferase (i.e. not associated to a restriction enzyme), some of which annotated specifically as *dam* (DNA adenine methylase) or *dcm* (DNA cytosine methylase). The largest plasmid, pKAZ3, carries as many as five methyltransferase genes. Secondly, other genes have annotations related to known anti-restriction functions. These include *ard* genes (alleviation of restriction of DNA), *ardA, ardB* and *ardC*, which have been shown to block restriction by various mechanisms (50); as well as *klcA* genes which are homologues of *ardB*; and the *vcrx089-093* operon which has recently been shown to provide anti-restriction and anti-CRISPR functions in *Vibrio* (29).

**Table 1.**
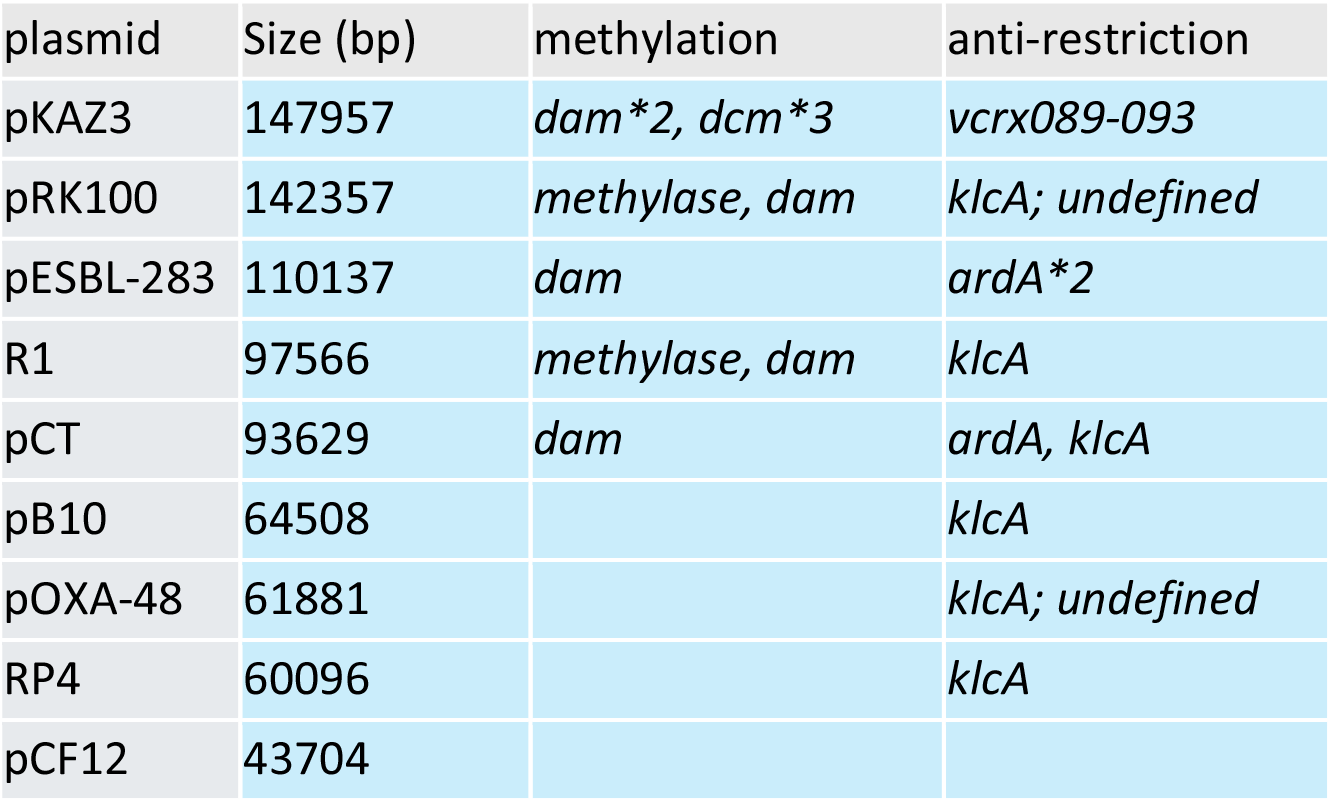
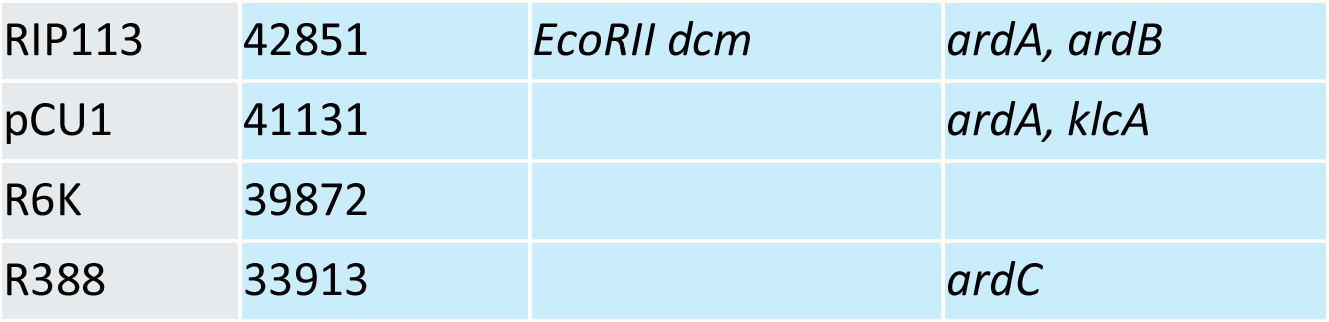
Summary of putative anti-restriction functions identified on plasmids. Plasmids are ordered from top to bottom by decreasing plasmid size. ‘undefined’ indicates genes for which ‘anti-restriction’ was the only information provided. Details are given in the text.

In addition to these plasmid functions, we noted that the chromosome of *E. coli* K-12 also carries methyltransferase and anti-restriction functions: this includes the well-known *dam* methyltransferase, *yhdJ* methyltransferase (which appears to have little effect without overexpression (51)), two copies of *klcA* in the cryptic prophages CP4-57 and CP4-6, and *dcm*. In particular, Dcm in *E. coli* K-12 strains is related to EcoRII methyltransferase, recognises the same sequence, and provides protection against cell killing linked to the loss of EcoRII RM system (52). This suggests that Dcm activity might lead to the absence of EcoRII restriction in our assays. We next analysed how a subset of these candidate genes affect plasmid transmission in the face of restriction.

### Effect of methyltransferases on restriction efficiency

We first asked if inactivating Dcm encoded on the *E. coli* K-12 chromosome allows the EcoRII RM system to be active against plasmids, by measuring plasmid conjugation for all wildtype plasmids, with and without Dcm in the donor and EcoRII RM system in the recipient. For most plasmids, the only combination displaying reduced conjugation was the combination of a *dcm-* donor with a *EcoRII+* recipient (Fig. S2), as expected if EcoRII restricts sites unprotected by Dcm methylation. However, interestingly RIP113 plasmid displayed lower conjugation efficiency from the *dcm-* donor even towards the *EcoRII-* recipient, and EcoRII had no additional effect on transfer efficiency (Fig. S2). Subsequent analysis of the RIP113 sequence revealed that it carries its own version of the EcoRII RM system (Table 1), explaining why it is protected from EcoRII restriction. Across plasmids, when plasmids were conjugated from the *dcm-* donor, the number of EcoRII recognition sites became a significant factor explaining EcoRII restriction efficiency (Fig. 3A, log_10_ (RE) ∼ log_10_ (n_sites_), F_1,38_ = 10.54, *p* = 0.0024), similarly to other Type II RM systems (Fig. 2). As before, this was not the case when using the *dcm+* donor (Fig. 3A, F_1,38_ = 2.08, *p* = 0.158). Still, when focusing on individual plasmids, we can see that 3 plasmids did not become susceptible to EcoRII, explaining why the effect across plasmids is relatively weak compared to other Type II RM systems. Among these 3 plasmids (pCU1, RIP113, pKAZ3), RIP113 carries EcoRII RM system, and pKAZ3 encodes 3 *dcm* genes, which might also contribute to methylation of EcoRII recognition sites.

We next tested if plasmid-encoded methyltransferases can also provide protection from restriction in our dataset, choosing pKAZ3 as an example as it contains both predicted adenine- and cytosine-methyltransferase genes (Table 1). We obtained knock-out mutants of one *dam* and one *dcm* gene, and compared restriction efficiency of each mutant with wild-type pKAZ3 (Fig. 3B). Surprisingly, pKAZ3 Δ*dam* was overall slightly less restricted than wild-type pKAZ3 across RM systems (2.1-fold increase, F_1,89_ = 11.0, *p* = 0.0013). However, there was no specific effect on a particular RM system (interaction F_9,80_ = 0.455, *p* = 0.9; all individual effects *p* > 0.05). On the other hand, for pKAZ3 Δ*dcm* there was a significant interaction between mutant status and RM system (F_9,136_ = 22.5, *p* < 2.10^-16^); by far the strongest effect was on restriction by EcoRII (t_14_ = 18.5, *p* = 3.18 10^-11^ after Bonferroni correction), pKAZ3 Δ*dcm* being restricted ∼ 2700-fold more than wild-type pKAZ3. In addition, *dcm* had a small but significant protective effect against EcoBI and EcoRV (Fig. 3B, respectively t_14_ = 3.38, *p* = 0.044 and t_14_ = 3.44, *p* = 0.04, after Bonferroni correction). Thus, we found that the *dam* gene we tested has no protective effect against restriction here, but *dcm* had a large protective effect against EcoRII, and a smaller effect against two other RM systems.

### Effect of other anti-restriction functions on restriction efficiency

Finally, we tested for an effect of other plasmid-encoded putative anti-restriction functions on restriction with our panel of RM systems. We focused again on pKAZ3, which carries an anti-defence operon which has been relatively less studied in comparison to better-known *ard* genes. The *vcrx089-vcrx093* operon was shown recently to have anti-restriction and anti-CRISPR activity in *Vibrio cholerae* in another IncC plasmid (29). Genes *vcrx089* and *vcrx090* had anti-restriction activity against *V. cholerae*’s Type I system, and genes *vcrx091-093* form a recombination system related to phage λ’s Red system, which repairs double-strand breaks due to CRISPR-Cas targeting (29). We hypothesized that these genes might also provide anti-restriction activity in *E. coli*, and tested this hypothesis by constructing two pKAZ3 variants, one with both *vcrx089* and *vcrx090* deleted, and one with a deletion of *vcrx093* (shown to be essential for anti-CRISPR activity in (29)).

We compared RE for wild-type, Δ*vcrx089-90* and Δ*vxrx093* pKAZ3 variants (Fig. 4). The knock-outs affected restriction overall, in a way that was dependent on the RM system tested (interaction effect, F_18,204_ = 5.88, *p* = 2.71 10^-11^). Each of the deletions led to increased sensitivity to specific RM systems, with variable amplitude. The Δ*vcrx089-90* deletion strongly increased RE by the related EcoBI and EcoKI systems (424-fold increase, t_14_ = 9.01, *p* = 6.6 10^-6^ and 906-fold increase, t_14_ = 10.9, *p* = 1.29 10^-6^ respectively) and had a smaller effect on EcoRV and EcoVIII RE (9.6-fold increase, t_14_ = 6.41, *p* = 3.24 10^-4^ and 12.2-fold increase, t_14_ = 11, *p* = 5.64 10^-7^ respectively). The Δ*vxrx093* deletion also had a large effect on EcoBI RE (426-fold increase, t_14_ = 6.76, *p* = 1.82 10^-4^), and smaller but significant effects on EcoAI, EcoRII and EcoRV restriction (respectively 13.1-fold increase, t_14_ = 3.74, *p* = 0.044; 5.67-fold increase, t_14_ = 3.85, *p* = 0.035; and 22.1-fold increase, t_14_ = 7.13, p=10^-4^). Thus, both *vcrx089-90* and *vxrx093* confer protection against restriction by various Type I and Type II RM systems.

Finally, we tested other genes with putative anti-restriction or repair activity (*ardB* and *umuCD* on RIP113, and *klcA* and *klcB* on pB10 plasmid), however we did not detect any significant effect of these gene deletions (p > 0.05 for all, Fig. S3).

## Discussion

In this work, we tested the activity of naturally occurring RM systems against a collection of conjugative plasmids that can transmit horizontally within *E. coli*. We found a large variation in restriction efficiency, which depends on the specific plasmid-RM pair; and uncovered some drivers of this variation.

First, restriction efficiency correlates with the number of RM recognition sites across conjugative plasmids. When analysed separately for each RM system, the correlation is significant mostly for Type II RM systems. This is in agreement with data obtained using phage λ mutants and EcoRI Type II system, in which the probability of phage escape from restriction decreased with each additional restriction site, and effects were multiplicative (53). This pattern is consistent with a simple mechanism of escape from restriction due to a low probability of stochastic methylation at each recognition site, and independent between sites. This also suggests that for Type II systems, additional plasmid-specific factors like anti-restriction functions have little role overall.

By contrast, for several Type I RM systems, we observe no significant correlation between restriction efficiency and recognition site number, suggesting that other factors drive the observed patterns. Indeed, we find many putative anti-restriction functions encoded on plasmids, particularly on the larger plasmids, in accordance with bioinformatic studies (26). We then show experimentally that some of these genes provide protection against specific RM systems. First, both host- and plasmid-encoded methyltransferases provide some protection. The effect of donor host methyltransferases on conjugation is well-known as a factor that can ameliorate the transmission of engineered conjugative vectors across species (54, 55). Here, we show that the chromosomal cytosine methylase Dcm protects all conjugative plasmids tested from restriction by EcoRII. This is likely to be more than an artefact from using a laboratory strain, as Dcm is widespread in *E. coli* natural isolates (56). Yet, some plasmids are still resistant to EcoRII when conjugating from a *dcm* mutant, and we show that in the case of pKAZ3, this is due to the presence of a plasmid-encoded cytosine methyltransferase. In contrast, we find no effect of inactivating a pKAZ3 adenine methyltransferase, but that plasmid contains two *dam* genes, likely providing functional redundancy. Adenine methylation was shown to provide resistance to restriction in another model conjugative plasmid (28). Still, methyltransferases could also have roles unrelated to restriction, including regulatory roles. For RIP113 plasmid, we observed a 10-fold decrease in conjugation even in the absence of restriction, when using a *dcm* mutant donor, suggesting that methylation contributes to the regulation of transfer, as shown for other conjugative plasmids (57).

In addition to methyltransferases, plasmids also carry other genes with putative anti-restriction (“ard”) functions. Known *ard* genes are predominantly active against Type I restriction, possibly because Type II systems are too diverse in terms of molecular structure and mechanism for a generic anti-restriction mechanism to evolve (25). Even within Type I systems, they are often active against only a subset of RM systems (58–60). In our assays, we detected no effects of *klcA* in pB10, or *ardB* in RIP113. *klcA* and *ardB* are related genes, and their activity *in vivo* has been shown to depend on their expression levels (61), explaining why *klcA* was previously observed to have no effect on restriction in RP4 plasmid (62). Instead, we find specific anti-restriction effect of other genes on pKAZ3 plasmid, for which an anti-defence role was demonstrated more recently in another IncC plasmid, pVCR94 (29). We find that *vcr089-vcr090* genes are active against some Type I as well as Type II RM systems, but quantitatively much more effective against Type I systems. We then demonstrate an anti-restriction role for gene *vcr093*, extending results from (29) which found a role against CRISPR-Cas interference. The anti-restriction mechanism is very likely the same as for its anti-CRISPR effect, namely by double-strand DNA break repair (29). Interestingly, this *vcrx089-093* anti-defence operon is not annotated as such by the annotation tools we used, and we only noticed it by using PaperBLAST (63). This strongly suggests that other anti-restriction functions on these plasmids still await detection.

Overall, our data shows that anti-restriction functions significantly contribute to the relative resistance of plasmids against RM systems, especially Type I systems, with 100-to 1000-fold increase in susceptibility to some Type I systems in anti-restriction gene knockouts. Alternatively, for some other plasmid-RM pairs, low plasmid sensitivity to restriction arises from carrying a low number of restriction sites, especially for Type II RM systems (Fig. 2). Interestingly, for several of the plasmids analysed here, the number of restriction sites is lower than expected by chance (Fig. S4): for instance, RP4, R388 and RIP113 would be expected to have 7, 5 and 9 EcoRI restriction sites respectively, based on their size and GC content, but they only have one. This is consistent with bioinformatic studies showing a pattern of Type II RM target avoidance in mobile plasmids, particularly in smaller plasmids (26). In small plasmids, the total number of recognition sites is lower, meaning that a mutation removing a single target site can have a large effect on restriction. Indeed, experiments on RP4 plasmid showed that adding three EcoRI sites led to a 10^4^ -fold increase in restriction (30). Finally, some general trends in our data could be explained by core features of conjugation. To contrast our results to measures of restriction against phages (the other main type of MGEs infecting bacteria), we compared our results to data from the BASEL collection of *E. coli* phages (32), which included the measure of restriction efficiency by six RM systems, four of which we tested against plasmids. We can discern two interesting patterns (Fig. S5). First, many phages are totally resistant to restriction, despite containing a large number of restriction sites. This is likely due to hypermodification of their genomes, as most of these phages belong to the Tevenvirinae family, which display cytosine hypermodification (a strategy that to our current knowledge is not available to plasmids). Secondly, plasmids have low sensitivity to restriction by Type III systems compared to many phages. This pattern is apparent across plasmids, suggesting it is not caused by plasmid-specific anti-restriction functions, but might instead be linked to conjugation itself. Indeed, experiments in the naturally transformable species *Neisseria gonorrhoeae* showed a 10^4^ -fold decrease in restriction efficiency for plasmid conjugation compared to transformation. This effect was attributed to restriction endonucleases acting on double-stranded (ds) DNA and not single-stranded (ss) DNA (64). Type III restriction requires the recognition of two unmodified recognition sites in inverse orientation (65, 66). Moreover, the endonuclease and methyltransferase sub-units are part of the same complex, with the Mod subunit providing specificity. Type III EcoPI was shown to methylate ssDNA *in vitro* (67). Thus, the dynamics of restriction and methylation on incoming plasmids might make conjugative entry intrinsically less susceptible to Type III restriction, similarly to filamentous, ssDNA phages which are also relatively resistant to Type III restriction (68). More generally, several factors point to a key role for the early dynamics of host defences and plasmid anti-defences and establishment functions in determining if conjugation is successful. Protective functions are expressed early during conjugation (69), and anti-defence genes are enriched in the leading region of plasmids (the first region entering the recipient during conjugation), suggesting that the speed of early events is critical (70).

Overall, our results show that restriction is on average low, and depends on the number of recognition sites on the plasmid (in particular for Type II restriction), and on the complement of anti-restriction functions carried by the plasmid (in particular for Type I restriction). The collection of plasmids we tested includes most types of plasmids carrying AMR genes in Enterobacteriaceae, in particular plasmids of groups IncF, IncI and IncN (5, 35), suggesting that these patterns might hold true for most clinically relevant AMR plasmids. In terms of RM systems, we only tested a limited subset of the large diversity in existing RM subtypes. Still, Type I and Type III systems are very similar in functional mechanism within Type, as most variation between systems is variation in site recognition by the specificity determinant (*hsdS* for Type I, *mod* for Type III) (71, 72), suggesting our results might be representative. For Type II systems, there is much more variation in mechanism, thus it might be difficult to establish general rules and other subtypes will need to be tested to establish generality.

Our results suggest that the overall effect of RM systems on plasmids in natural populations will be relatively low, due to multiple plasmid strategies mitigating restriction. Still, even a relatively poor defence system should have relevant effects on plasmid and bacterial eco-evolutionary dynamics. For a given plasmid, even a two-fold restriction barrier would lead to a large competitive disadvantage in competition with another plasmid unaffected by restriction. On the host side, RM system distribution will determine which bacteria preferentially receive a plasmid in mixed populations. Restriction against a costly plasmid should provide a fitness benefit, while restriction against a beneficial plasmid should be counter-selected (73). Moreover, RM distribution might also impact selective pressures acting on the donor: investment in conjugative transmission is costly for the donor bacteria, so is not directly selected for. However, because clonemates share the same RM systems, RM will bias transfer preferentially to bacterial clonemates (40). In turn, this can allow increased investment by donors to transmit beneficial plasmids, via kin selection (74), because the benefits of transfer will be borne preferentially by the donors’ kin. Overall, these direct and indirect effects of RM-plasmid interactions will combine with the (likely stronger) effects of defence against phage infection (31) to impact RM distribution across bacteria. Finally, bacteria in nature commonly carry multiple defence systems, including on average more than two RM systems (14). The next step will thus be to study not only clinically relevant plasmids, as we have done here, but also clinically relevant and genetically more complex bacterial hosts (3), to understand patterns of AMR plasmid spread in clinically-relevant bacterial communities (75).

## Data availability

The data underlying this article are available in the article and in its online supplementary material. Sequences for plasmids pCU1 and RIP113 are available in the European Nucleotide Archive, project PRJEB76555.

## Acknowledgments

We thank Calin Guet for sharing RM-system carrying bacterial strains, and Allison Lopatkin and Jeronimo Rodriguez-Beltran for sharing conjugative plasmids.

