## Supplementary Figures and Tables for "Various plasmid strategies limit the effect of bacterial Restriction-Modification systems against conjugation"

| plasmid | size (bp) | MOB | MPF | Replicon | AMR genes | antibiotic for selection | reference sequence |
| --- | --- | --- | --- | --- | --- | --- | --- |
| R388 | 33913 | - | - | IncW | AAC(6')-Ib7;dfrB2;qacEdelta1;sul1 | trimethoprim | NC_028464 |
| R6K | 39872 | MOBP | typeT | IncX2 | APH(3'')-Ib;APH(6)-Id;TEM-181 | streptomycin | LT827129.1 |
| pCU1 | 41131 | MOBF | typeT | IncN | AAC(6')-Ib7;ANT(3'')-IIa;OXA-2;OXA-2;qacEdelta1;sul1 | streptomycin |  |
| RIP113 | 42161 | MOBF | typeT | IncN | tet(A) | tetracycline |  |
| pCF12 | 43704 | MOBP | typeT | IncX3 | QnrS1;SHV-66 | ampicillin | MT720906 |
| RK2 | 60096 | MOBP | typeT | IncP1 | APH(3')-Ib;TEM-181;tet(A) | tetracycline | BN000925.1 |
| pOXA48 | 61881 | MOBP | typeI | IncL/M | OXA-48 | ampicillin | NC_019154.1 |
| pB10 | 64508 | MOBP | typeT | IncP1 | AAC(6')-Ib7;APH(3'')-Ib;APH(6)-Id;OXA-2;qacEdelta1;sul1;tet(A) | tetracycline | AJ564903.1 |
| pCT | 93629 | MOBP | typeI | IncB/O/K/Z | CTX-M-14 | ampicillin | FN868832.1 |
| R1 | 97566 | MOBF | typeF | IncFII | ANT(3'')-IIa;APH(3')-Ia;TEM-181;catI;qacEdelta1;sul1 | streptomycin | NZ_KY749247.1 |
| pESBL283 | 110137 | MOBP | typeI | IncI1 | AAC(6')-Ib7;ANT(3'')-IIa;CTX-M-1;dfrA1;qacEdelta1;sul1;sul2 | streptomycin | NC_024978.1 |
| pRK100 | 142357 | MOBF | typeF | IncFIB;IncFII | TEM-181;tet(A) | tetracycline | NZ_CP060383.1 |
| pKAZ3 | 147957 | MOBH | typeF | IncA/C2 | AAC(6')-Ib7;QnrVC1;VEB-9;dfrA1;dfrA23;qacEdelta1;qacEdelta1;sul1;sul1;tet(A);tet(C) | tetracycline | NZ_MT720905.1 |

**Table S1: wildtype conjugative plasmids used in this study.**

| **RM system** | **type** | **strain** | **plasmid** | **recognition sequence** (5’- -3’) |
| --- | --- | --- | --- | --- |
| EcoAI | IB | NK354 | NA | GAGNNNNNNNGTCA |
| EcoBI | IA | WA251 | NA | TGANNNNNNNNTGCT |
| EcoKI | IA | MG1655 | NA | AACNNNNNNGTGC |
| EcoR124I | IC | NK402 | NA | GAANNNNNNRTCG |
| EcoRI | IIP | MG1655 ΔhsdS::KnR | pMB1 ori, bla | GAATTC |
| EcoRII | IIEP | MG1655 ΔhsdS::KnR | pMB1 ori, bla | CCWGG |
| EcoRV | IIP | MG1655 ΔhsdS::KnR | pMB1 ori, bla | GATATC |
| EcoVIII | IIP | MG1655 ΔhsdS::KnR | pMB1 ori, bla | AAGCTT |
| EcoP1 | III | MG1655 ΔhsdS::KnR | p15A ori, cat | AGACC |
| EcoP15I | III | MG1655 ΔhsdS::KnR | p15A ori, cat | CAGCAG |

**Table S2: RM systems used in this study.** R stands for A or G; W stands for A or T.

| number | | Name | Sequence |
| --- | --- | --- | --- |
| TD_RM1 | H1_dcm_p1 | | TGTAATTATGTTAACCTGTCGGCCATCTCAGATGGCCGGTGAAATCTATGGTGTAGGCTGGAGCTGCTTC |
| TD_RM2 | H2_dcm_p2 | | TCGCGCGCATATTTTTGCTGCGAGTGGCCTTATCGTGAACGTCGGCCATGATGGGAATTAGCCATGGTCC |
| TD_RM3 | dcm - 1,854,543 F | | TTTTCATCCGCTGGCTGAG |
| TD_RM4 | dcm - 1,856,105 R | | CGCTTCTCTATCGCCGTATC |
| TD_RM5 | H1_pB10_klcA_p1 | | CTTGAAGGCCGGGCATTTCGCCCAGGTCAACTACCGGAGAAAGTCCGATGGTGTAGGCTGGAGCTGCTTC |
| TD_RM6 | H2_pB10_klcA_p2 | | GCCCCGGCCGAAGCCGGGGCCGCCCCTCATCAGTCAATAGCCCGGTAGATATGGGAATTAGCCATGGTCC |
| TD_RM7 | H1_pB10_klcB_p1 | | CGCGACGTGATCGCGTGGCGGCGTGCTGACACTTGAGGGGCCGGCCGATGGTGTAGGCTGGAGCTGCTTC |
| TD_RM8 | H2_pB10_klcB_p2 | | CGTCGTTCGTCATTCGTTCGTTCTCCTGGTCAAGTGATAAGCCGCAACTGATGGGAATTAGCCATGGTCC |
| TD_RM9 | pB10_klcA - 40,987 F | | GTTGTTCCTGGTCATGGTCG |
| TD_RM10 | pB10_klcA - 41,599 R | | AGGGAAAGGGCAAGGTTTAG |
| TD_RM11 | pB10_klcB - 39,515 F | | CACTCCAGCCGGATATTGG |
| TD_RM12 | pB10_klcB - 40,868 R | | GGTTCAGCTCGGGATTCTG |
| TD_RM13 | H1_pKAZ3_dam1_p1 | | TTCGTTTCTACCTCTTGAATGTTTCGTCAGAACAAACTGGAGCGCTTATGGTGTAGGCTGGAGCTGCTTC |
| TD_RM14 | H2_pKAZ3_dam1_p2 | | ATGCAAAACCCGCCAATCGGCGGGTTTTTTTATGCTGCCACGGTAGCGGCATGGGAATTAGCCATGGTCC |
| TD_RM15 | pKAZ3_dam1 - 22,794 F | | AGGTCTTCGTTCTTGAACCG |
| TD_RM16 | pKAZ3_dam1 - 23,647 R | | TGCCCCGACTTTATCAAGAC |
| TD_RM17 | H1_pKAZ3_dcm3_p1 | | TTCCCGATTGGGACGGTAGTGGCCTCTCAACCGAAAGGAGAGAGCTCATGGTGTAGGCTGGAGCTGCTTC |
| TD_RM18 | H2_pKAZ3_dcm3 p2 | | AGGCCTTTCGGCATTGCCCCGGTGAGTTACTAATGAACCAAAGATGAAGTATGGGAATTAGCCATGGTCC |
| TD_RM19 | pKAZ3_dcm3 - 80,458 F | | ACGACTCTGACGAAACGAAG |
| TD_RM20 | pKAZ3_dcm3- 82,422 R | | CTGGCAATGCAGTGTTGAAG |
| TD_RM21 | H1_vcrx089 | | CCGCCCGGTTCGGGACGTGGTGCGCCTGTCACAATAGGAGCGCACCTATGGTGTAGGCTGGAGCTGCTTC |
| TD_RM22 | H2_Vxrx090 | | CCCTTTCGGGGAGCAAGCCCCCGTAAGGGTTAAAAAAAGTGATCCTCTTGATGGGAATTAGCCATGGTCC |
| TD_RM23 | H1_Vcrx093 | | CTCCCCCGCTGGGGGAGATCTCCCTCCTAACCATTGGAGAGAGGTCCATGGTGTAGGCTGGAGCTGCTTC |
| TD_RM24 | H2_Vcrx093 | | CCAAGGGAAAGCGGGGGAAAATCCCCCACTCACCAGTAGAACGTCTCTACATGGGAATTAGCCATGGTCC |
| TD_RM25 | pKAZ3 - 66,670 F vcrx08990 | | TCTTCGGAGAGCGAAAAGTG |
| TD_RM26 | pKAZ3 - 68,800 R vcrx08990 | | CAGAACCGAGATTACCGACC |
| TD_RM27 | pKAZ3_dvcrx093F | | CCTTCGTGACAAGGAGATGG |
| TD_RM28 | pKAZ3_dvcrx093R | | TGTTCACCCTTGTTACACCG |
| TD_RM29 | H1_ardB_RIP113 | | TACCGCCGCCCGAAAAATTGCCGGTATCGGCTTAAAGAGGAAGTATCATGGTGTAGGCTGGAGCTGCTTC |
| TD_RM30 | H2_arcB_RIP113 | | AAGGCGGGCTTTGCCCGCCTGGTTATCAGTTAATCAATGGCACGATAAATATGGGAATTAGCCATGGTCC |
| TD_RM31 | RIP113_ardB - 1,123 F | | CAGACGGTAGGGTGTTGAAG |
| TD_RM32 | RIP113_ardB - 1,781 R | | CTTCCATGCCCTGACTGG |
| TD_RM33 | H1_RIP113_umuC | | CCATAACAGCGATACTGTATAAATAAACAGTTATTTGGAAGATCGCTATGGTGTAGGCTGGAGCTGCTTC |
| TD_RM34 | H2_RIP113_umuD | | GTTAACCGGCCCGGGTACGGCGCCGGTAATTATTTGATGGTGGCTATTGGATGGGAATTAGCCATGGTCC |
| TD_RM35 | RIP113umuC - 1,934 F | | ATATAGCGTTCGTTGGCAGC |
| TD_RM36 | RIP113umuD - 3,808 R | | GAAACCGTGCCGAAACTTG |

**Table S3: primers used in this study.**

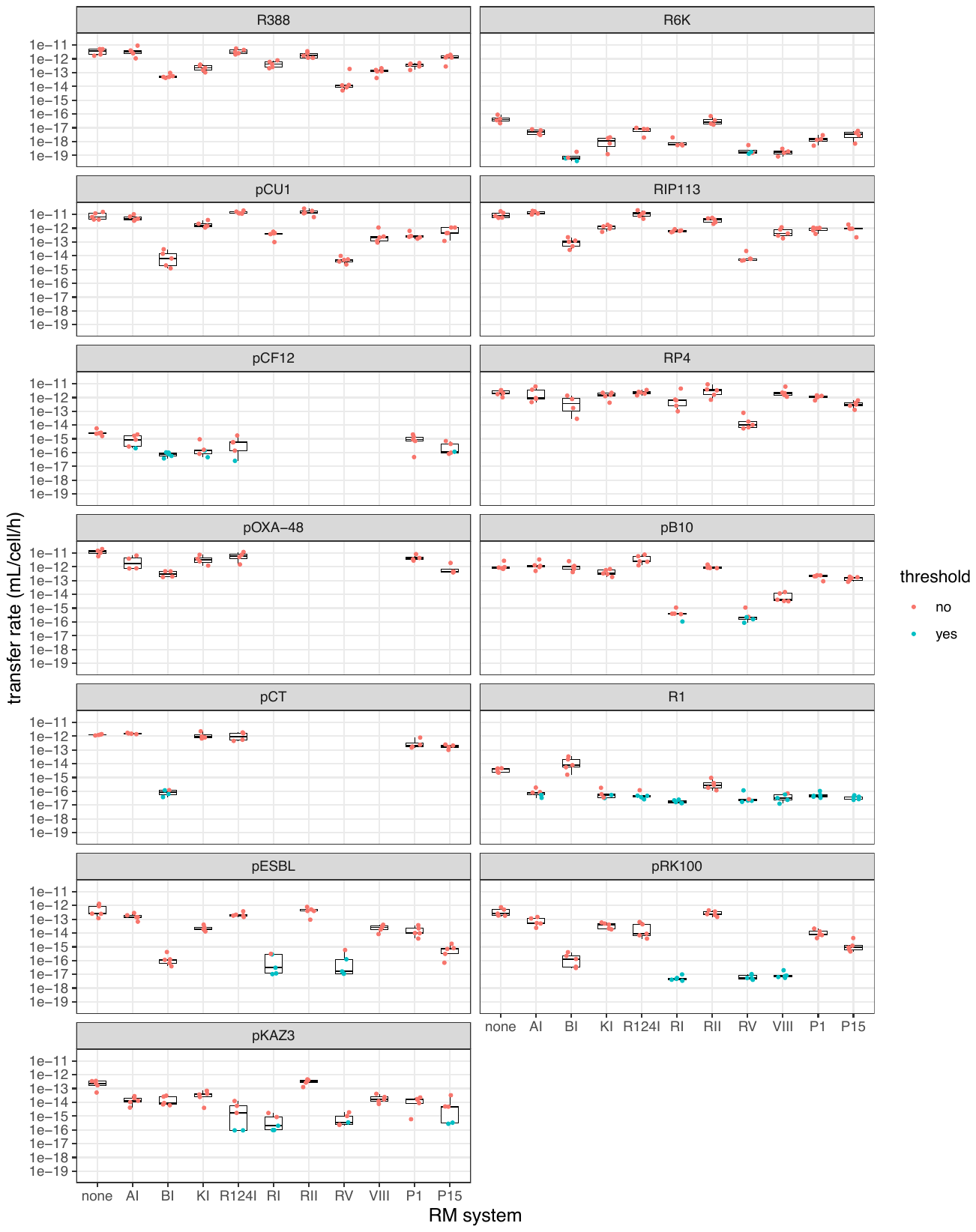

**Figure S1: transfer efficiency of wildtype plasmids.** Each facet shows transfer efficiency for one of 13 wildtype plasmids, as a function of the *E. coli* RM system present in the recipient strain. The centre value of the boxplots shows the median, boxes the first and third quartile, and whiskers represent 1.5 times the interquartile range. Individual replicates are shown as dots (n≥4); red dots indicate that transconjugant colonies were counted, whereas blue dots indicate that no transconjugants were observed and the threshold transfer efficiency is shown instead.

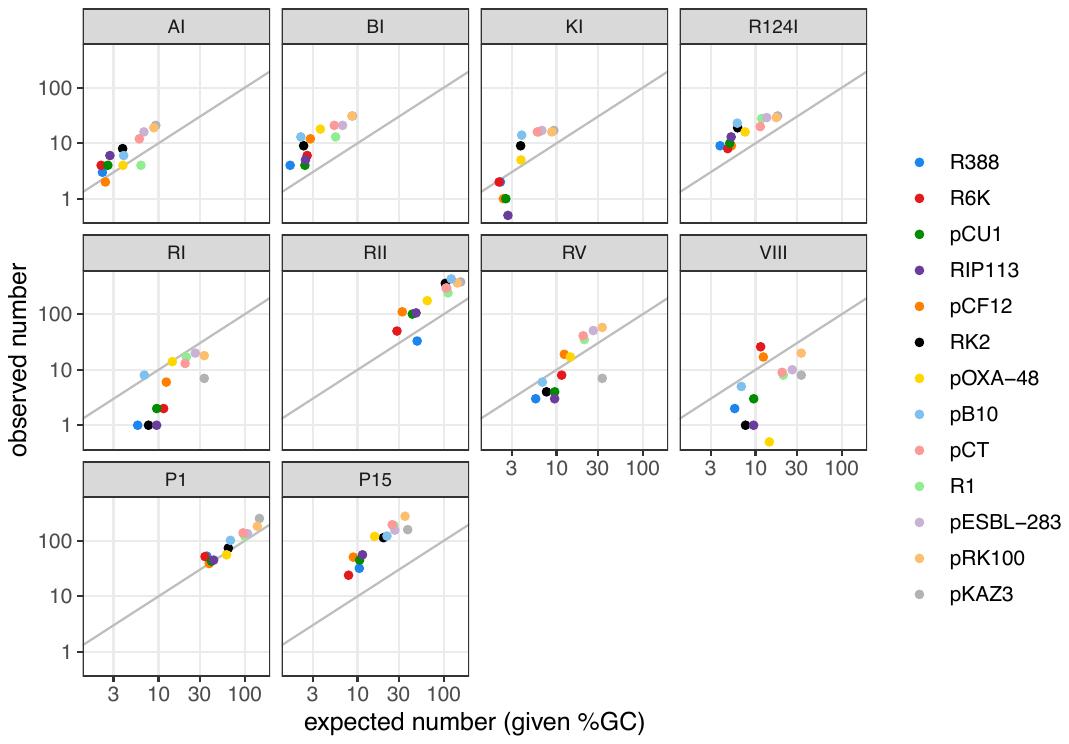

**Figure S2: expected and measured number of RM recognition sites**, ordered by RM system. The expected number of recognition sites is calculated by taking into account the base composition of the recognition sequence, and the overall %GC of the plasmid considered, using the following formula: 0.5 (1-%GC)^n( A+T)^ * 0.5 (%GC)^n( G+C)^*sequence length. Observed values are shown as a function of expected values, as a log scale.

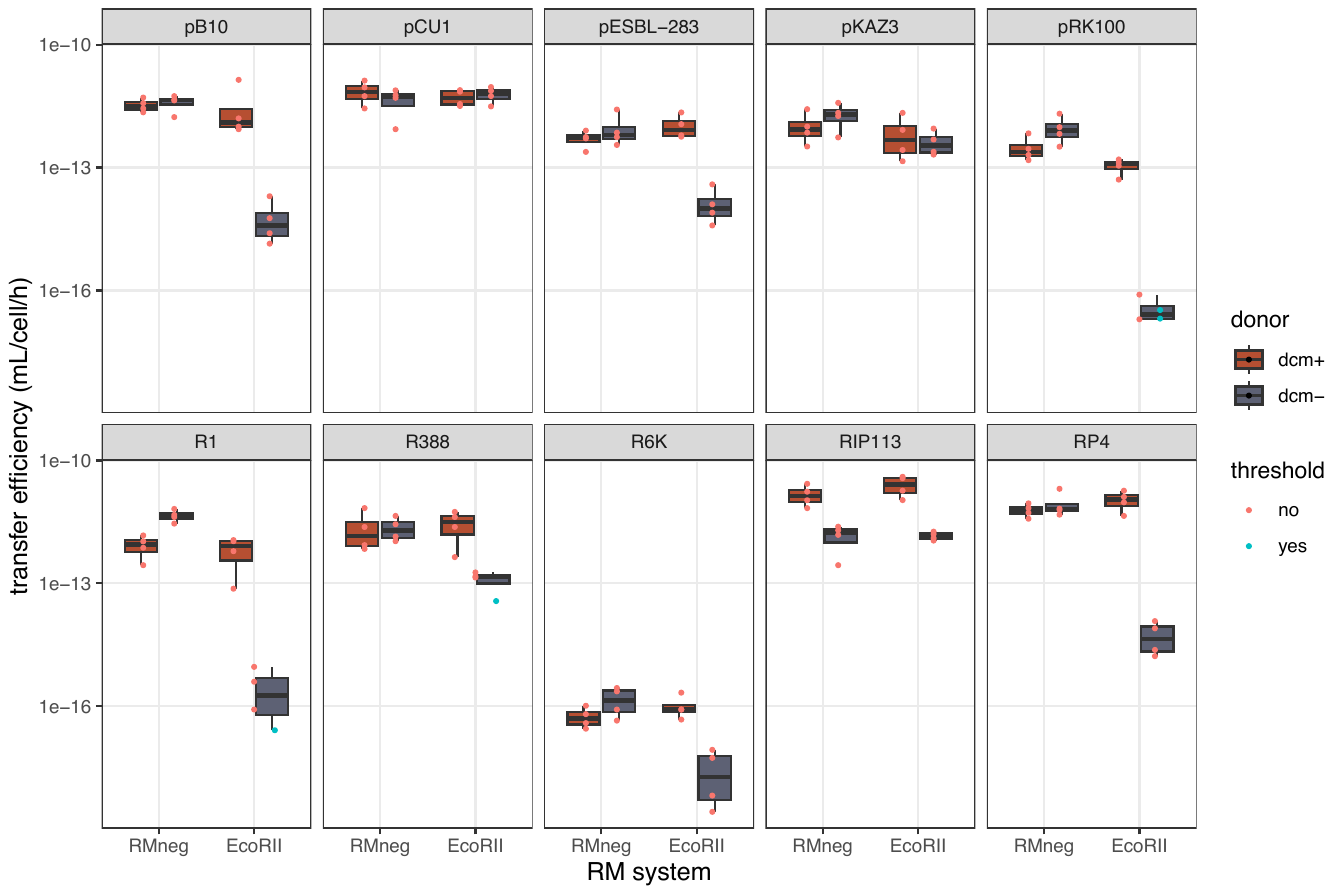

**Figure S3: transfer efficiency of wildtype plasmids from *dcm+* and *dcm-* donors.** Each facet shows transfer efficiency for one of 10 wildtype plasmids, as a function of the *E. coli* RM system present in the recipient strain (none or EcoRII). The centre value of the boxplots shows the median, boxes the first and third quartile, and whiskers represent 1.5 times the interquartile range. Individual replicates (n = 4) are shown as dots; red dots indicate that transconjugant colonies were counted, whereas blue dots indicate that no transconjugants were observed and the threshold transfer efficiency is shown instead.

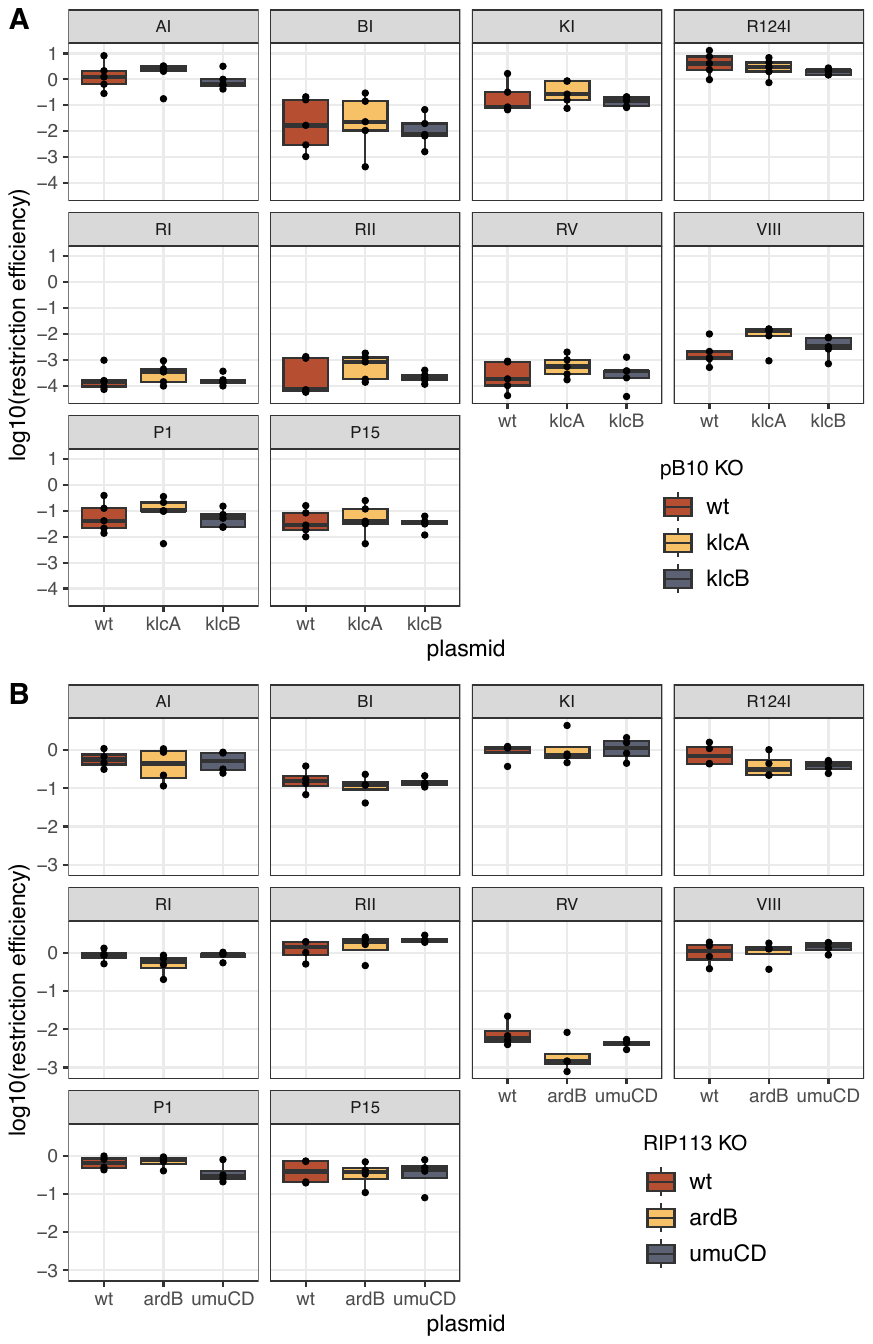

**Figure S4: effect of additional plasmid anti-restriction functions on restriction efficiency.** Log-transformed restriction efficiency for 10 *E. coli* RM systems is shown from a *dcm+* donor strain for wild-type and mutant pB10 (A) and wild-type and mutant RIP113 (B). The centre value of the boxplots shows the median, boxes the first and third quartile, and whiskers represent 1.5 times the interquartile range. Individual replicates are shown as dots (n≥4). Mutant plasmids showed no significant difference from wild-type plasmids in restriction efficiency in any RM treatment after Bonferroni correction (all *p* > 0.05).

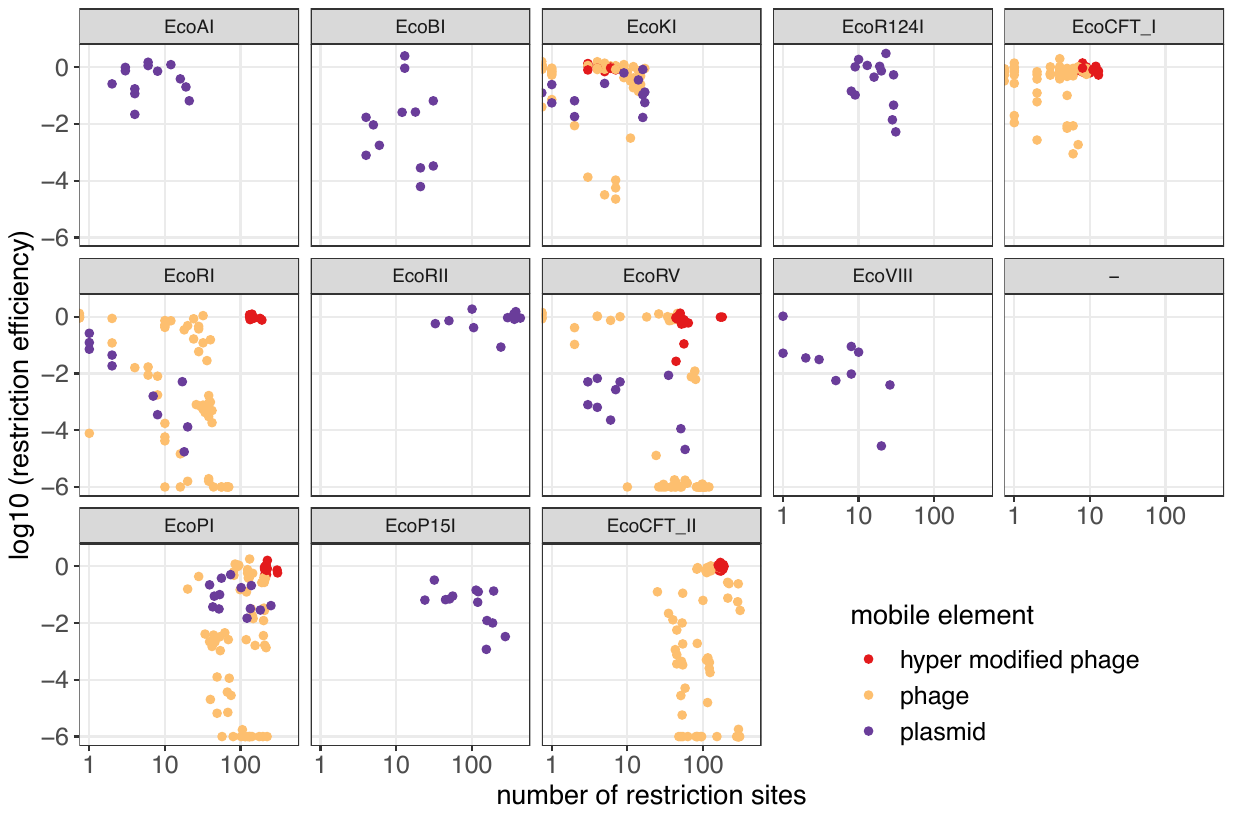

**Figure S5: comparison of restriction efficiency against conjugative plasmids and phages.** Mean values of restriction efficiency data presented in Fig. 2 for conjugative plasmids and mean values of restriction efficiency calculated from EOP (efficiency of plating) data in ^[1]^ (Maffei et al.) using the BASEL collection of *E. coli* phages, are shown as a function of RM recognition site number on each element. Plasmid data are shown in violet; phage data are shown in orange, except phages from the Tevenvirinae (T-even phages), which are known to be hyper-modified and shown in red. A value of 10^-6^ was added to the phage EOP data as an estimated detection threshold limit similar to the threshold applied to our conjugation data.

1. Maffei et al., ‘Systematic Exploration of Escherichia Coli Phage–Host Interactions with the BASEL Phage Collection’.
